# Reemployment of Kupffer’s vesicle cells into axial and paraxial mesoderm via transdifferentiation

**DOI:** 10.1101/2021.11.29.470501

**Authors:** Takafumi Ikeda, Kiichi Inamori, Toru Kawanishi, Hiroyuki Takeda

**Affiliations:** Laboratory of Embryology, Department of Biological Sciences, Graduate School of Science, The University of Tokyo, 7-3-1 Hongo, Bunkyo-ku, Tokyo 113-0033, Japan

**Keywords:** charon, zebrafish, Kupffer’s vesicle, transdifferentiation

## Abstract

Kupffer’s vesicle (KV) in the teleost embryo is a fluid-filled vesicle surrounded by a layer of epithelial cells with rotating primary cilia. KV transiently acts as the left-right organizer but degenerates after the establishment of left-right asymmetric gene expression. Previous labelling experiments indicated that descendants of KV-epithelial cells are incorporated into mesodermal tissues after KV collapses (KV-collapse) in zebrafish embryos. However, the overall picture of their differentiation potency had been unclear due to the lack of suitable genetic tools and molecular analyses. In the present study, we established a novel zebrafish transgenic line with a promoter of *charon*, in which all KV-epithelial cells and their descendants are specifically labelled until the larval stage. We found that KV-epithelial cells underwent epithelial-mesenchymal transition upon KV-collapse and infiltrate into adjacent mesodermal progenitors, the presomitic mesoderm and chordoneural hinge. Once incorporated, the descendants of KV-epithelial cells expressed distinct mesodermal differentiation markers and contributed to the mature populations such as the axial muscles and notochordal sheath through normal developmental process. These results indicate that fully differentiated KV-epithelial cells possess unique plasticity in that they are reemployed into mesodermal lineages through transdifferentiation after they complete their initial role in KV.

## Introduction

Kupffer’s vesicle (KV) is a teleost-specific organ, which is transiently formed on the ventral side of the embryonic tailbud (Brummett and Dumont, 1978; Kupffer, 1868). KV consists of a fluid-filled lumen and its surrounding epithelial cells (hereafter referred to as KV-epithelial cells), and functions as the left-right organizer (LRO), playing a crucial role in left-right axis formation in the embryo (Essner et al., 2005). In the zebrafish, KV-epithelial cells are derived from a special group of cells called dorsal forerunner cells (DFCs; Melby et al., 1996; Oteíza et al., 2008). DFCs originally bear endodermal character as they express endoderm-specific genes, *sox32* and *sox17* (Alexander et al., 1999; Kikuchi et al., 2001; Warga and Kane, 2018). During epiboly stages, DFCs are transformed into KV-epithelial cells through mesenchymal-epithelial transition (Amack et al., 2007; Matsui et al., 2015; Oteíza et al., 2008; Zhang et al., 2016). At early somite stages, each KV-epithelial cell protrudes a primary cilium on the apical side, which rotates and generates the leftward fluid flow in the KV lumen (Essner et al., 2005). Sensed by primary cilia themselves, the leftward flow induces left-right asymmetric gene expression in and around KV, leading to left-sided expression of *nodal* in the lateral plate mesoderm (LPM; Essner et al., 2005). Judged by their morphological characteristics and specialized function, KV-epithelial cells are considered as fully differentiated and functional cells.

In the light of the fate plasticity in matured cells, we were interested in the fate of KV-epithelial cells after they complete their mission in left-right patterning. Indeed, KV collapses and disappears soon after the establishment of the asymmetric gene expression in the LPM (hereafter referred to as KV-collapse), but previous lineage-tracing studies of DFCs demonstrated that upon KV-collapse, descendants of DFCs are incorporated into mesodermal tissues such as notochord, somites and tail mesenchyme (Cooper and D’Amico, 1996; Melby et al., 1996). However, detailed tracking and molecular characterization of those incorporated cells have not been performed so far, and thus we do not know yet whether ciliated KV-epithelial cells actually possess fate plasticity to transdifferentiate into functional mesodermal cells.

Recently, several transgenic zebrafish lines that express fluorescent proteins in DFCs and KV were generated for easier and more reliable tracing of DFC- and KV-derived cells. Among them, *sox17* lines have been frequently used because *sox17* promoter uniformly labels both DFCs and KV-epithelial cells from epiboly to somitogenesis stages (Compagnon et al., 2014; Mizoguchi et al., 2008; Oteíza et al., 2008; Sakaguchi et al., 2006). Furthermore, a recent work using *sox17:GFP-CAAX* line reported the epithelial-to-mesenchymal transition (EMT) of KV-epithelial cells during KV-collapse (Amack, 2021), but their fate after EMT has not been directly tracked yet. For specific fate tracking of KV-epithelial cells, the *sox17* lines are not necessarily suitable, because endodermal cells as well as DFCs are broadly labelled (Mizoguchi et al., 2008). Other transgenic lines marking KV-epithelial cells (summarized in Table 1) have a similar limitation (Caron et al., 2012; Chen et al., 2012; Du and Dienhart, 2001; Molina et al., 2007). Thus, for precise tracing, another transgenic line was required in which mature KV-epithelial cells are specifically and uniformly labelled.

**Table 1:**
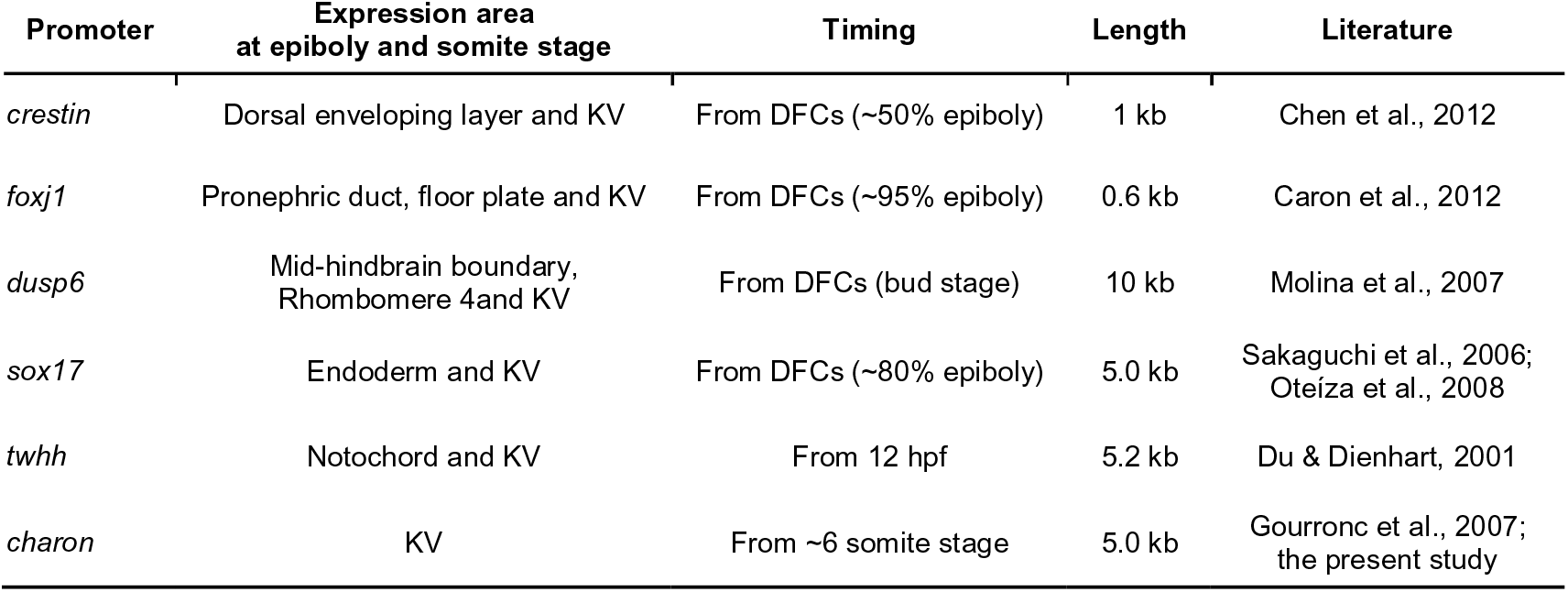
Transgenic lines in zebrafish for the visualization of DFCs and KV-epithelial cells

Here, we established novel transgenic lines, in which entire KV-epithelial cells are specifically labelled, using the promoter of *charon*. *charon* encodes a secreted Nodal antagonist belonging to the DAN family (Hashimoto et al., 2004). Its expression starts shortly after the differentiation of KV-epithelial cells from DFCs, and is highly specific to KV-epithelial cells both in zebrafish and medaka (Hashimoto et al., 2004; Hojo et al., 2007). The expression of *charon* is initially symmetric, but gradually becomes right-sided under the control of the leftward flow in KV (Hojo et al., 2007; Lopes et al., 2010). The essential role of *charon* in the left-right pattering is conserved among vertebrates as *charon*-knockout mice and zebrafish both exhibit randomized left-right axis (Marques et al., 2004; Montague et al., 2018).

Using these *charon* lines, we first confirmed that after KV-collapse, KV-derived cells (KVDCs) differentiate into mesodermal but not into ectodermal or endodermal tissues. Detailed timelapse imaging and gene-expression analyses further revealed that KVDCs undergo EMT and infiltrate into two types of undifferentiated mesoderm, which are the presomitic mesoderm and the chordoneural hinge. Importantly, they expressed differentiation markers similar to their surrounding mesodermal cells at later stages. These findings demonstrate that KV-epithelial cells possess high plasticity after completing their initial role in KV, and that they transdifferentiate into mesodermal lineages during normal development.

## Results

### Establishment of *charon:EGFP* to specifically label KV-epithelial cells

To establish transgenic lines in which KV-epithelial cells are specifically labelled, we cloned the 5-kb upstream sequence of zebrafish *charon* (Fig.1a, Supplementary Fig.1). A previous transient promoter assay demonstrated that the upstream sequence of zebrafish *charon* drives the reporter expression around KV (Gourronc et al., 2007), but any stable transgenic line using the *charon* promoter has yet to be established. After confirming the previous report by the Tol2-mediated EGFP reporter assay (Supplementary Fig.3a), we established three stable transgenic lines in which either EGFP, Lck-mGreenLantern (mGL) or H2B-mEosEM (a nuclear-localizing form of a photoconvertible fluorescent protein mEosEM; Fu et al., 2020) expression is driven by the *charon* 5-kb promoter (Fig.1b, Supplementary Fig.4a). In all the lines established, reporter gene expression was specifically detected in KV-epithelial cells (Fig.1c-e, Supplementary Fig.4b, c). In *charon:EGFP* embryos, EGFP fluorescence started to be detected from around the 6-somite stage (ss) just like endogenous *charon* (Hashimoto et al., 2004), and became stronger as KV grows in size. DAB (3,3’-Diaminobenzidine, tetrahydrochloride) immunohistochemistry against EGFP showed that at 6 ss and 12 ss, EGFP expression is strictly limited to the entire KV-epithelial cells with no ectopic expression in adjacent mesoderm (Fig.1d-f). Fluorescence *in situ* hybridization with immunofluorescence (FISH-IF; He et al., 2020) of *charon* in *charon:EGFP* embryos further showed that *charon* expression mostly overlaps with that of EGFP at 12 ss (Fig.1g). However, unlike endogenous *charon*, EGFP expression did not exhibited left-right bias (Fig.1d-f). This can be explained by the fact that 3’ untranslated sequence of *dand5* (a mouse ortholog of *charon*) is responsible for its right-sided expression (Minegishi et al., 2021; Nakamura et al., 2012). Untranslated sequences of zebrafish *charon*, which are not included in the plasmid construct of this study, could have a similar function to that of mouse *dand5*.

**Figure 1.**
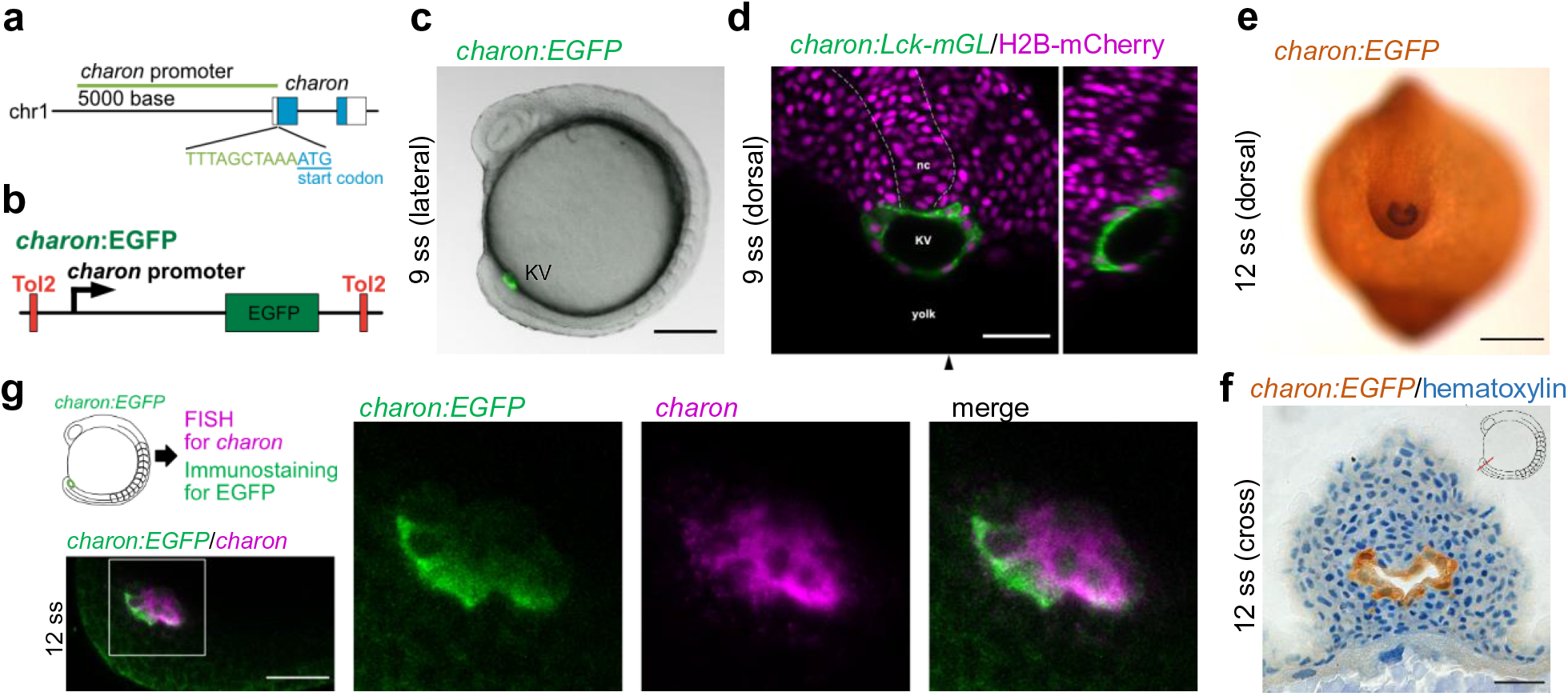
Establishment of *charon:EGFP* for the specific labeling of KV-epithelial cells **a**, Location of the 5-kb *charon* promoter in the zebrafish genome. *charon* resides on chromosome 1, and the 5-kb *charon* promoter spans from the direct upstream position of its start codon. **b**, Construction of *charon:EGFP* transgene flanked by Tol2 excision sites. **c**, Lateral view of the EGFP fluorescence in a *charon:EGFP* embryo at 9 somite stage (ss). Fluorescence image was merged with the DIC (differential interference contrast) image. KV, Kupffer’s vesicle. **d**, Dorsal view of a *charon:Lck-mGL* embryo at 9 ss injected with H2B-mCherry mRNA. Black arrowhead indicates the level of the sagittal optical section on the right side. **e**, Dorsal view of a DAB-stained *charon:EGFP* embryo at 12 ss. **f**, Cross section of the tailbud of a DAB-stained *charon:EGFP* embryo at 12 ss. Nuclei were counterstained with hematoxylin. Dorsal side to the top. **g**, FISH-IF of *charon* in a *charon:EGFP* embryo at 12 ss. Magenta signal is for FISH of *charon* mRNA whereas green is for immunostaining of EGFP proteins. Lateral view of the tailbud region is shown. White box indicates the enlarged area. Dorsal to the bottom, posterior to the left. Scale bars, 200 µm (c, e), 100 µm (f) and 50 µm (d, h).

Furthermore, we found that the activity of upstream sequences of *charon* in KV-epithelial cells is conserved among teleosts. A transgenic medaka (*Olyzias latipes*), which was established with the upstream sequence of medaka *charon* (Supplementary Fig.2), exhibited EGFP-reporter expression in KV-epithelial cells from 4 ss with faint ectopic expression in the notochord (Supplementary Fig.5c). Overall, these results show that *charon* transgenic lines are best suitable for the study of development and function of KV-epithelial cells.

### KVDCs differentiate into mesodermal lineages

KV is a transient organ and starts to degenerate at mid-somite stages (14–16 ss) when asymmetric *nodal* expression is established in the LPM (Cooper and D’Amico, 1996; Long, 2003; Supplementary Fig.8). However, we observed that EGFP-expressing cells in *charon:EGFP* embryos survived after KV-collapse and even persisted in hatched larvae (Supplementary Fig.3). Hereafter we refer to the EGFP-positive cells in *charon:EGFP* after KV-collapse as “KV-epithelium-derived cells (KVDCs)” to distinguish them from KV-epithelial cells. To track KVDCs after KV-collapse, we histologically analyzed embryos at the mid-somite stage (22 ss) and larvae at 3 days post fertilization (dpf) using DAB immunohistochemistry and confocal microscopy. In 22-ss embryos whose KVs have already diminished, most KVDCs were clustered around the collapsed lumen, but some of them were found in the adjacent notochord and presomitic mesoderm (PSM; Fig.2a, b). In 3-dpf hatched larvae, KVDCs were broadly distributed in the region posterior to the cloaca where no endodermal tissue exists (Fig.2c). KVDCs were detected in mesodermal tissues including both segmented and unsegmented posterior-most somites (Moriyama et al., 2012; Fig.2c, e1, e2), notochord cells near the posterior end and the notochordal sheath in the more anterior side (Fig.2d, e3), and fin mesenchymal cells with radial filopodia especially in the caudal fin fold (Fig.2c, d and e4). We counted the number of KVDCs in each tissues using *charon:H2B-mEosEM* 3-dpf larvae and found that they were most abundant in the unsegmented posterior-most somites (56±11 cells/embryo; Fig.2f). In contrast, no KVDCs were observed in the ectoderm (Fig.2d, e).

**Figure 2.**
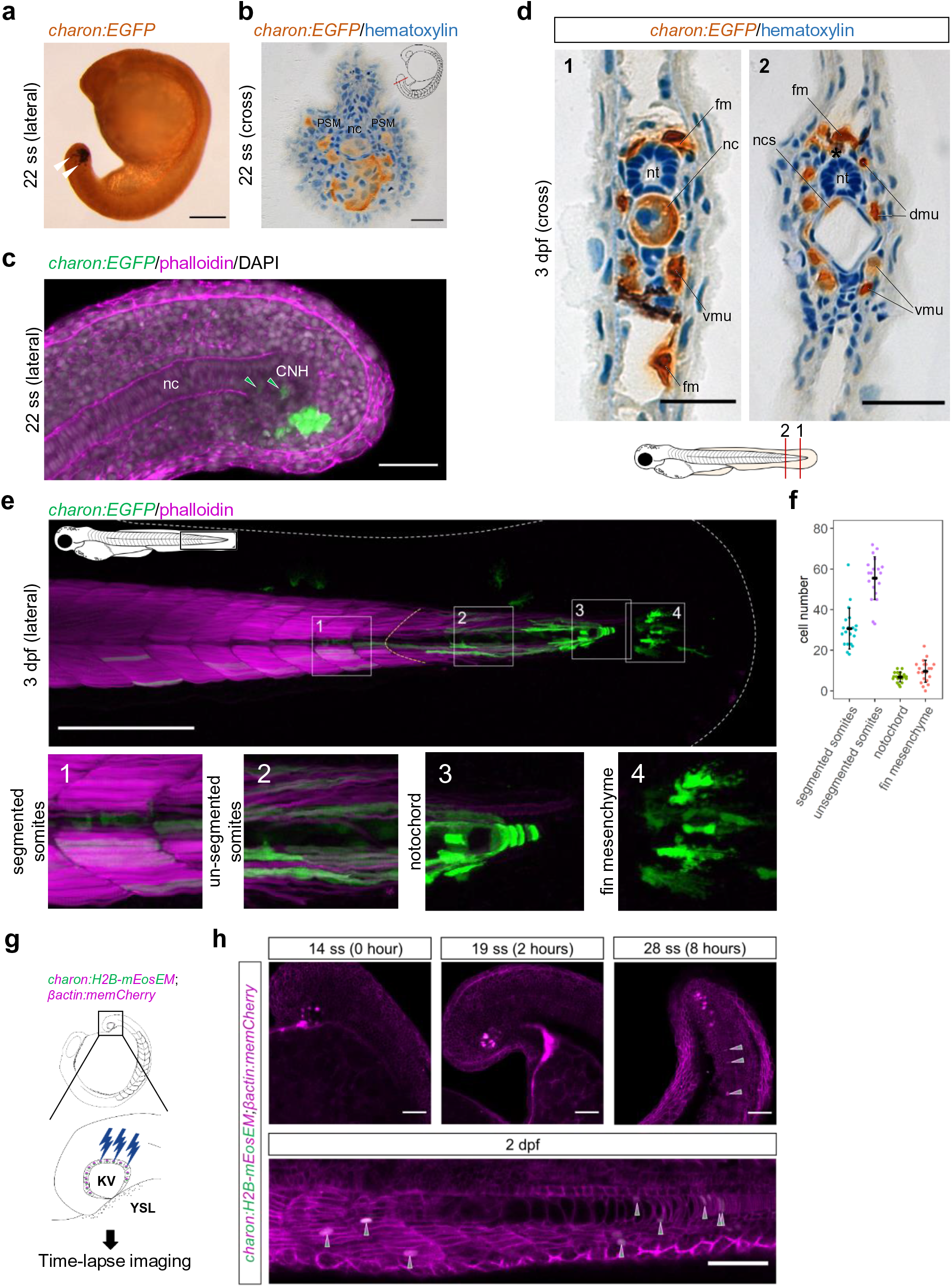
Histology of KVDCs **a**, Lateral view of a DAB-stained *charon:EGFP* embryo at 22 ss. White arrowheads indicate KVDCs which have started to migrate. **b**, Cross section of the tailbud of a DAB-stained *charon:EGFP* embryo at 22 ss. Nuclei were counterstained with hematoxylin. Dorsal side to the top. PSM, presomitic mesoderm. nc, notochord. **c**, Lateral view of a *charon:EGFP* embryo at 22 ss counterstained with phalloidin and DAPI. Dorsal to the top. Green arrowheads indicate EGFP-positive cells in the CNH (chordoneural hinge). **d**, Cross sections of a DAB-stained *charon:EGFP* larva at 3 dpf. Nuclei were counterstained with hematoxylin. Dorsal side to the top. Asterisk indicates a melanocyte. dmu, dorsal muscle. fm, fin mesenchyme. nc, notochord. ncs, notochordal sheath. nt, neural tube. vmc, ventral muscle. **e**, Lateral view of the caudal region of a *charon:EGFP* embryo at 3 dpf counterstained with phalloidin. White boxes indicate areas magnified in the panels 1-4. White dashed line indicates the edge of the fin fold. Panel 1, EGFP-positive muscles in segmented somites. Panel 2, EGFP-positive muscles in non-segmented somites. Panel 3, EGFP-positive notochord cells. Panel 4, EGFP-positive fin mesenchymal cells in the caudal fin fold. Scale bars, 200 µm (a, e), 100 µm (b, d), 50 µm (c). **f**, Number of H2B-mEosEM-positive KVDCs in *charon:H2B-mEosEM* in four populations shown in (e). Horizontal bars, mean. Vertical bars, standard deviation. N = 20 larvae. **g**, Scheme for the labelling of KV-epithelial cells by photoconversion. H2B-mEosEM is expressed in KV-epithelial cells in β*actin:memCherry* embryos by injection of *charon:H2B-mEosEM* plasmid. H2B-mEosEM was photoconverted at around 13 ss using a confocal microscopy and time-lapse images were taken every 1 hour. **h**, Time-lapse images of a β*actin:memCherry* embryo injected with *charon:H2B-mEosEM*. Images taken every 2 hours after photoconversion and from 2 dpf treated larvae are shown. Arrowheads indicate the migrated KVDCs labelled with photoconverted H2B-mEosEM.

To further support the above findings, we directly examined the migration of KVDCs into these mesodermal tissues by a photoconversion-based method (Shimada et al., 2013). We photoconverted H2B-mEosEM transiently expressed in KV-epithelial cells of β*actin:memCherry* embryos, whose cell membrane is labelled with mCherry (Xiong et al., 2014). After photoconversion at around 13 ss, labelled KV-epithelial cells were tracked at an interval of 1 hour during segmentation periods (n = 4 embryos, Fig.2g). At 14 ss (0 hour after photobleaching), photoconverted H2B-mEosEM was only detected in KV-epithelial cells, confirming the specificity of photoconverted labelling (Fig.2h). The labelled KVDCs tended to stay together until 19 ss (4 hours after photobleaching), and then gradually migrated from KV into the PSM and notochord (Fig.2h, arrowheads in the 28-ss panel). When the same embryos were further observed at 2 dpf, the labelled KVDCs were broadly distributed in segmented somites, unsegmented posterior-most somites, and the notochord (Fig.2h, arrowheads in the 2-dpf panel). We obtained a similar result using another transgenic line, *foxj1a:KikGR*, in which photoconvertible protein KikGR (Tsutsui et al., 2005) is expressed in KV-epithelial cells as well as in the floor plate (Caron et al., 2012; Supplementary Fig.7a). Taken together, the results demonstrated that KVDCs migrate and contribute to the axial and paraxial mesoderm.

### Loss of the LRO and epithelial character in KVDCs

We next examined the molecular characteristics of KVDCs after KV-collapse. We first examined whether any KVDCs are eliminated by apoptosis during and after KV-collapse. Immunostaining of cleaved caspase-3, an executioner of apoptotic process (Elmore, 2007), revealed that apoptosis was induced in KVDCs after KV-collapse, but that the number of apoptotic KVDCs was limited (Fig.3a, b). We then asked how survived KVDCs lost morphological and molecular characteristics of the LRO components by examining *charon* (LRO marker) and *sox17* (DFCs marker) expression and the protein distribution of ZO-1 (epithelial marker) and acetylated α-tubulin (primary cilia marker) in KVDCs (Alexander and Stainier, 1999; Essner et al., 2005; Hashimoto et al., 2004; Oteíza et al., 2008). FISH-IF showed that the expression of *charon* was diminished at 20 ss and 26 ss compared to at 12 ss, and *charon*-negative KVDCs were frequently detected (Fig.3c). Moreover, we did not detect *sox17* expression in KV-epithelial cells or KVDCS at all stages examined (Supplementary Fig.6), which is consistent with the previous report (Alexander and Stainier, 1999).

**Figure 3.**
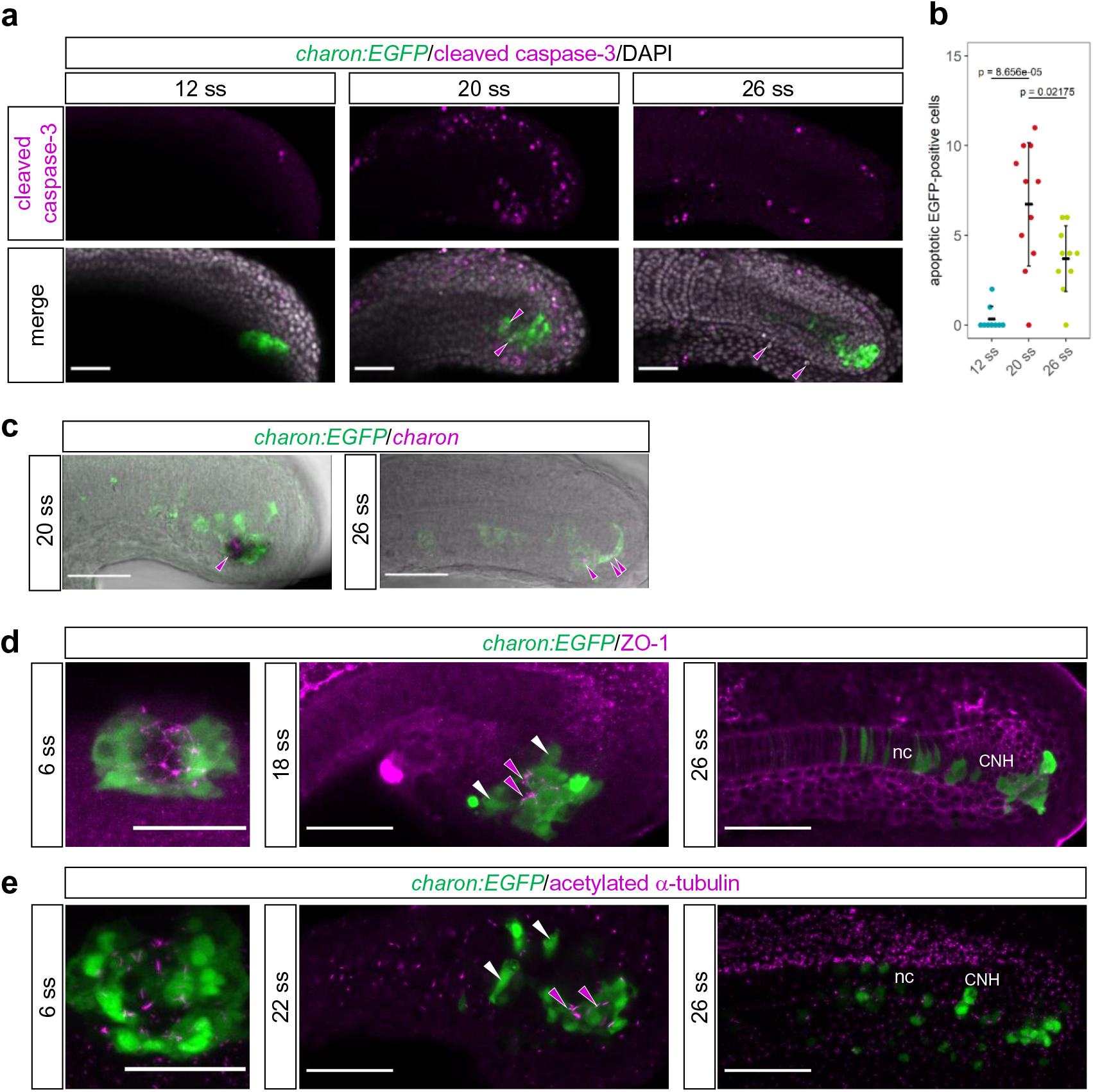
Loss of the LRO character in KVDCs **a**, Immunostaining of cleaved caspase-3 in *charon:EGFP* embryos at 12, 20 and 26 ss. Top, the signal of cleaved caspase-3. Bottom, merged view with EGFP and DAPI signal. Magenta arrowheads indicate KVDCs which is positive for cleaved caspase-3. **b**, Quantification of cleaved caspase-3-positive KVDCs. Horizontal bars, mean. Vertical bars, standard deviation. *p-*values from two-tailed Welch’s t-test are shown. N = 9 (12 ss), 11 (20 ss), and 10 (26 ss) embryos. **c**, FISH-IF of *charon* in *charon:EGFP* at 20 and 26 ss. Magenta, signal of FISH for *charon* mRNA. Green, signal of immunostaining for EGFP. Lateral view of the tailbud region is shown. Dorsal to the top. **d**, Immunostaining of ZO-1 in *charon:EGFP*. KV is shown for 6 ss whereas the tail region is shown for 18 and 26 ss. Magenta and white arrowheads indicate KVDCs with and without ZO-1 expression, respectively. nc, notochord. CNH, chordoneural hinge. **e**, Immunostaining of acetylated α-tubulin in *charon:EGFP*. Magenta and white arrowheads indicate KVDCs with and without acetylated α-tubulin signal, respectively. Scale bars, 50 µm (a, c, d, e).

ZO-1 was accumulated on the surface of the KV lumen at 6 ss, confirming the epithelial character of EGFP-positive cells in *charon:EGFP* (Fig.3d; Oteíza et al., 2008). At 18 ss, shortly after KV-collapse, migrating KVDCs no longer expressed ZO-1, although residual ZO-1 expression was detected in a cluster of KVDCs located on the ventral side of the CNH. At 26 ss, a substantial number of KVDCs moved to the posterior edge of the chordoneural hinge (CNH), a group of progenitors which are located at the posterior end of the notochord and thought to give rise to the notochord, floor plate and hypochord (Agathon et al., 2003; Row et al., 2015). Furthermore, primary cilia of KVDCs were largely diminished after KV-collapse, particularly in migrating ones (Fig.3e). Taken together, the results suggest that KV-epithelial cells gradually lose their LRO and epithelial character, and acquire mesodermal fates through mid-to late somite stages.

### KVDCs undergo EMT at the onset of migration

We examined detailed cellular behavior of KVDCs at the onset of migration using a double transgenic line, *charon:EGFP;*β*actin:memCherry*. KV initially protruded into the yolk, then incorporated into the tailbud at around 10 ss, and finally collapsed (Supplementary Figure 8). At around 14 ss, KV-epithelial cells detached from the underlying yolk syncytial layer (YSL) (Fig.4, Supplementary Video 1). Interestingly, a ring-like structure labelled by strong mCherry signals appeared in the YSL just beneath KV (Fig. 4, magenta arrowheads). This YSL ring shrank when KV was detached from the YSL as if it pushes KV out into the tailbud and breaks the connection between KV and the YSL. Once KV was detached from the YSL, its lumen started to shrink (Fig. 4, the 01:20 panel). At the same time, KV-epithelial cells started to disperse and migrate, extending filopodia-like structures (Fig. 4, insets in the 02:40 panel, Supplementary Video 2). This observation, together with the loss of ZO-1 expression (Fig.3d), suggests that KV-epithelial cells undergo epithelial-mesenchymal transition during KV-collapse.

**Figure 4.**
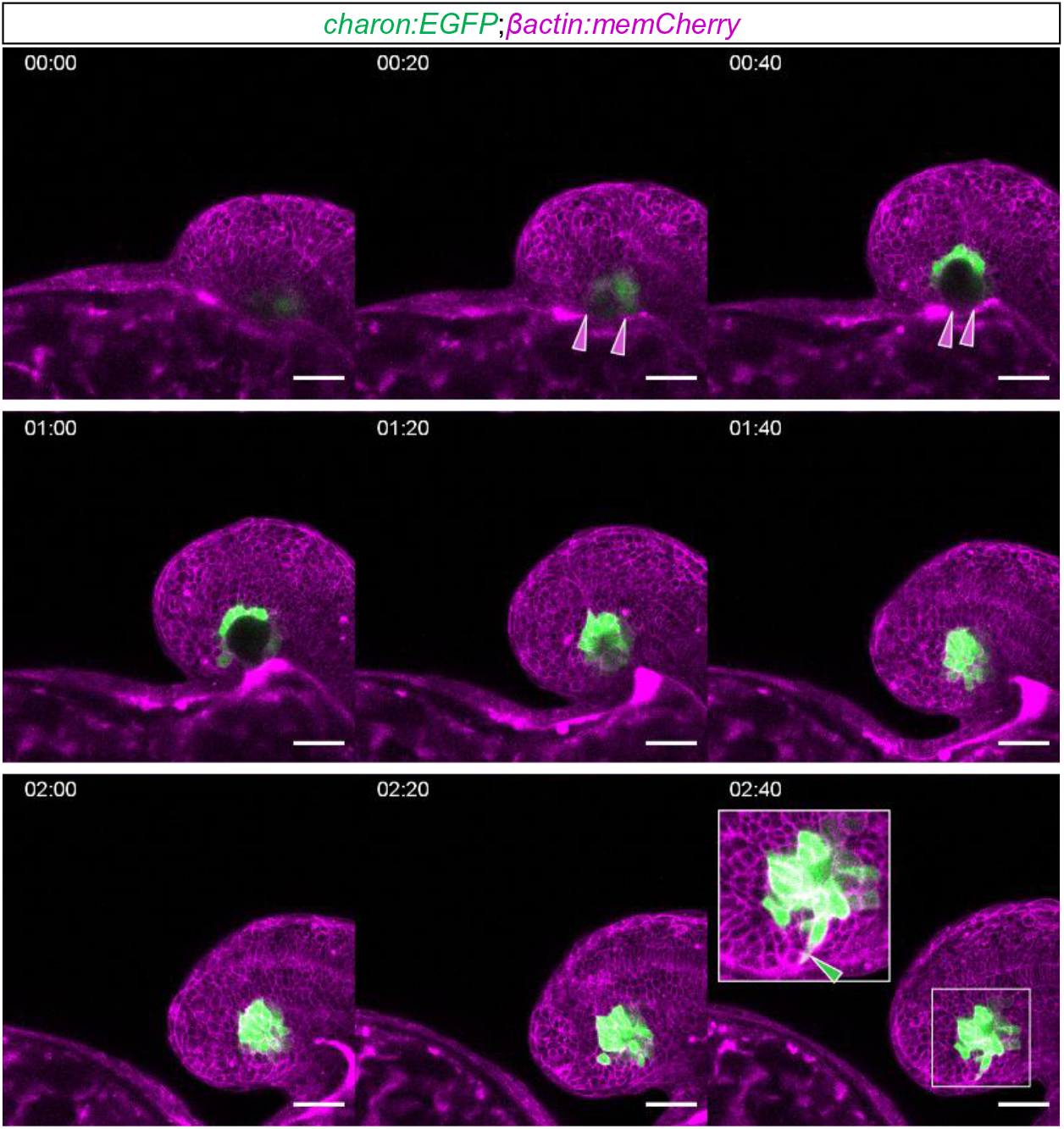
KVDCs migrate to mesodermal tissues through EMT Time-lapse images of a *charon:EGFP;*β*actin:memCherry* embryo taken every 20 minutes for 3 hours from 13 ss. Magenta arrowheads indicate the shrinkage of the YSL underlying KV. White boxes in the panel 02:40 indicate inset areas. Green arrowhead in the inset indicates a filopodium-like structure formed in KVDCs. Scale bars, 50 µm.

### Differentiation of KVDCs into the PSM and somite derivatives

We next examined the developmental capacity of KVDCs which migrate into somites using lineage-specific differentiation markers. At 12 ss when KV is still present, KV epithelial cells were ventrally attached to the PSM as well as notochord (Fig.5a). A pan-PSM marker *msgn1* (Joseph and Cassetta, 1999; Yoo et al., 2003; Yoon et al., 2000) was strongly expressed in the PSM but not in KV-epithelial cells at 12 ss (Fig.5a), confirming their distinct lineages. After KV-collapse, KVDCs were found in the PSM (Fig.5b1) and later in the anterior segmented somites as well, where *msgn1* expression is no longer detected (Fig.5b2). These results suggest that KVDCs are incorporated into the PSM after KV-collapse and are intermingled with surrounding PSM cells.

**Figure 5.**
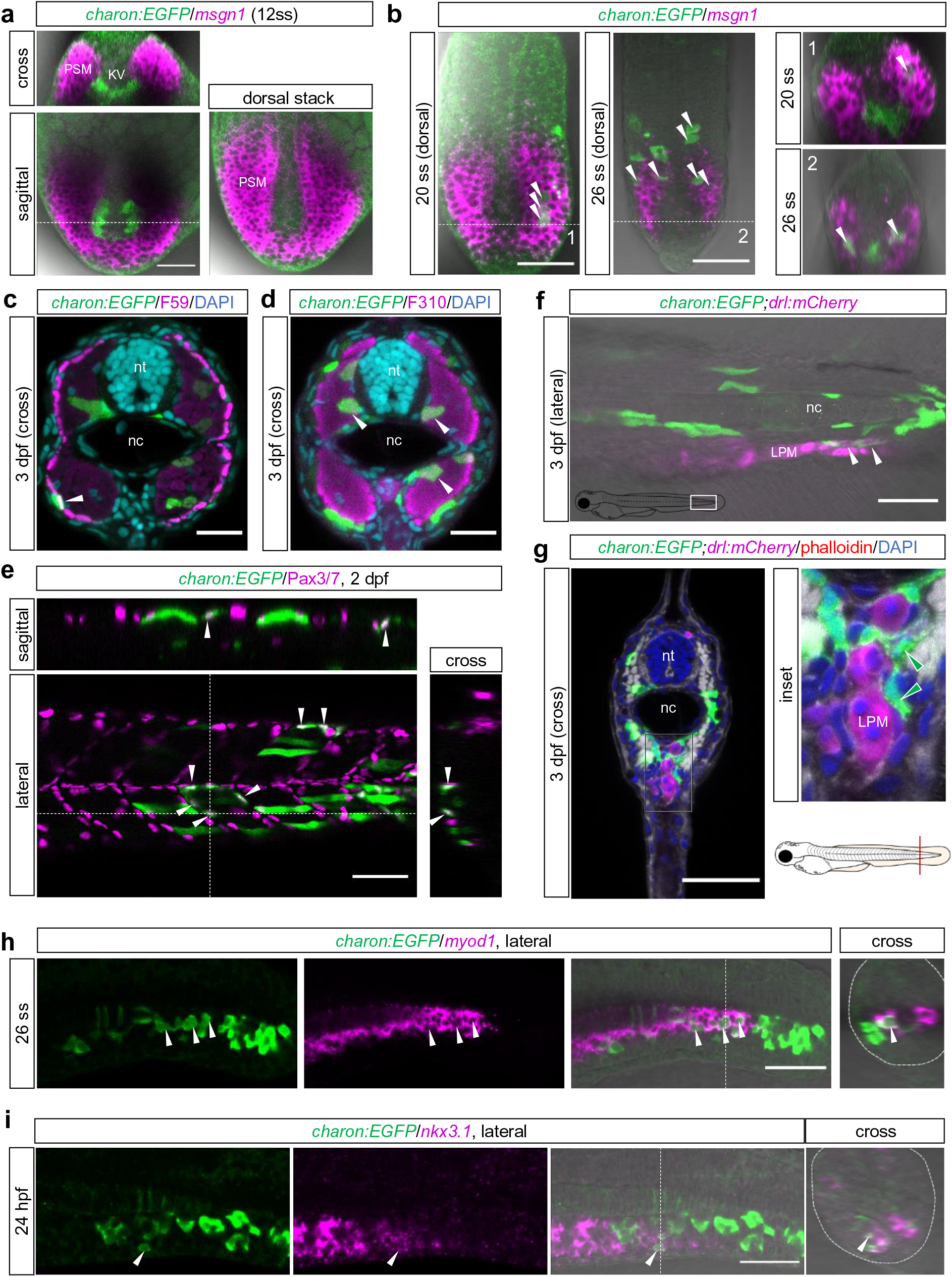
Differentiation of KVDCs into somite derivatives **a**, FISH-IF of *msgn1* in a *charon:EGFP* embryo at 12 ss. Left, dorsal view. Top, an optical section at the level of the dashed line in the panel below. Right, a more dorsal stack than the left one. PSM, presomitic mesoderm. **b**, FISH-IF of *msgn1* in *charon:EGFP* embryos at 20 ss and 26 ss. Left, dorsal view. Anterior to the top. Right, optical sections at the level of the dashed lines in the left panels. White arrowheads indicate KVDCs in the PSM (20 ss and 26 ss) and in the segmented somites (26 ss). **c, d**, Vibratome cross sections of *charon:EGFP* 3 dpf larvae immunostained with F59 (c) and F310 (d) antibodies to show the distribution of slow and fast muscles, respectively. Counterstaining was performed with DAPI. White arrowheads indicate EGFP-positive slow and fast muscle cells. Dorsal to the top. nt, neural tube. nc, notochord. **e**, Immunostaining of dermomyotomes by anti-Pax3/7 antibody in a *charon:EGFP* 2 dpf larva. Lateral view and optical sections at the level of dashed lines are shown. White arrowheads indicate EGFP-positive dermomyotomes. **f**, Lateral view of a *charon:EGFP;drl:mCherry* 3 dpf larva. White arrowheads indicate EGFP-positive sclerotomal cells surrounding mCherry-positive cells derived from the LPM. nc, notochord. LPM, lateral plate mesoderm. **g**, Left, cross section of a *charon:EGFP;drl:mCherry* 3 dpf larva immunostained with anti-EGFP and anti-mCherry antibodies. Counterstaining was performed with phalloidin and DAPI. Right, a magnified view of the white rectangular area in the left panel. White arrowheads indicate KVDCs surrounding mCherry-positive LPM derivatives. **h, i**, FISH-IF of *myod1* in a *charon:EGFP* embryo at 26 ss (h) and *nkx3.1* in a 24 hpf (hours post fertilization) embryo (i). Left panel, lateral view. Anterior to the left, dorsal to the top. Right panel, optical sections at the level of the dashed lines in the left panels. White arrowheads indicate KVDCs which differentiated in *myod1-*positive adaxial cells (h) and *nkx3.1*-positive sclerotomes (i). Scale bars, 50 µm (a-i).

The zebrafish somite consists of three populations with distinct fates, the myotome, dermomyotome, and sclerotome (Stickney et al., 2000). The myotome is further divided into the two subpopulations, slow muscle in the dorsal (outer), and fast muscle in the ventral (deeper) regions. KVDCs were found to be distributed in all these populations at later stages (Fig.5). At 3 dpf, KVDCs which are located near the surface and in the deep layer of the myotome expressed slow and fast muscle markers, respectively, as detected by F59 and F310 antibodies (Fig.5c, d). These muscle cells persisted at least until 5 dpf (data not shown), suggesting that they differentiated into functional muscles. Fast muscles are known to be derived from adaxial cells which are large, cuboidal in shape, located adjacent to the notochord and express *myoD* (Devoto et al., 1996). During slow muscle development, adaxial cells radially migrate from their original position toward the superficial layer of the myotome and differentiate into slow muscles there (Barresi et al., 2001; Cortés et al., 2003). At late somite stage (26 ss), we identified adaxial cell-like KVDCs, judged by their shape and location, and found that they indeed expressed *myoD* (Fig.5h). This observation suggests that KVDCs follow the normal developmental process to differentiate into slow muscles.

KVDCs were also found in the dermomyotome layer (the outer-most surface of the somite) and in the sclerotome region (the ventro-medial region near the notochord and neural tube), where they expressed the dermomyotome marker, Pax3/7 (Hammond et al., 2007) and sclerotome marker, *nkx3.1* (Ma et al., 2018), respectively (Fig.5e, i). We also examined the positional relationship between ventral KVDCs and posterior blood vessels, because the ventral sclerotome is known to differentiate into mural cells, which surround blood vessels (Stratman et al., 2017). For this, we used double transgenic fish of *charon:EGFP* and *drl:mCherry*, whose LPM derivates are specifically labelled with mCherry (Mosimann et al., 2015; Prummel et al., 2019). We found that phalloidin-negative KVDCs were attached to the posterior-most part of the caudal vein (Fig.5f, g), supporting the idea that KVDCs acquire the differentiation potency of the sclerotome. Importantly, KVDCs did not overlap with mCherry-expressing LPM-derivatives, indicating that they do not contribute to the LPM lineage (Fig.5f, g).

### Differentiation of KVDCs into the notochord and tail mesenchyme

We further examined the differentiation process of KVDCs in the notochord and tail mesenchyme. At 12 ss, KV-epithelial cells were ventrally attached to the *no tail* (*ntl*, the young notochord marker)-positive notochord (Schulte-Merker et al., 1994; Fig.6a). At 20 ss, some KVDCs were found in the posterior end of *ntl*-expressing area, i.e., the CNH (Fig.6b). These KVDCs were also positive for *ntl*, indicating that they had acquired the axial mesodermal fate. Some KVDCs further developed into *ntl-*negative mature notochord cells at 26 ss (Fig.6b), and later into the surrounding sheath cells (Fig.2d). These results show that KVDCs which infiltrate into the CNH possess the full capacity of notochord differentiation.

**Figure 6.**
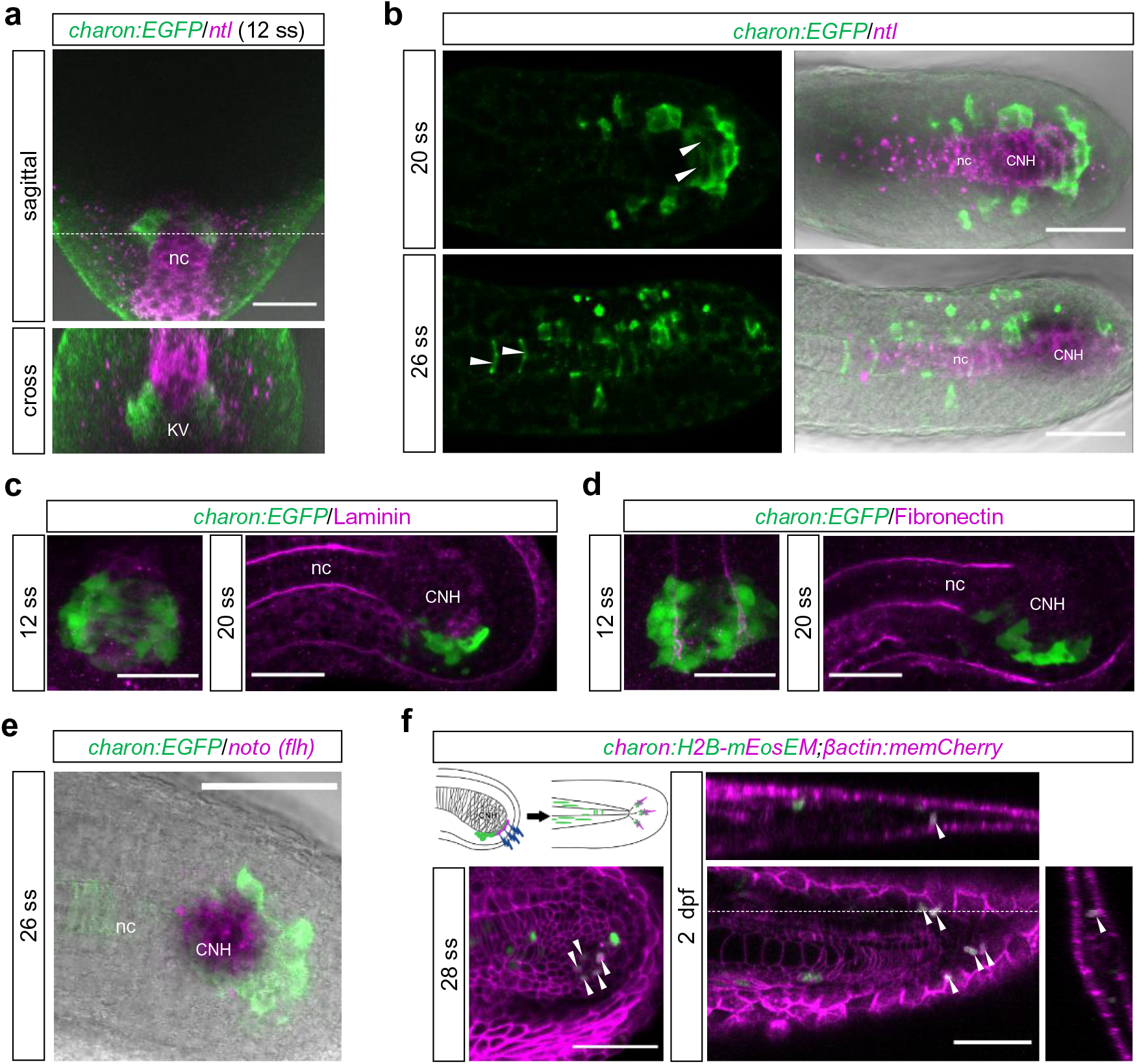
Differentiation of KVDCs into the notochord **a**, FISH-IF of *ntl* in a *charon:EGFP* embryo at 12 ss. Top, dorsal view. Anterior to the top. Bottom, an optical section at the level of the dashed line in the upper panel. nc, notochord. **b**, FISH-IF of *ntl* in *charon:EGFP* embryos at 20 ss and 26 ss. Dorsal views. Anterior to the left. White arrowheads, KVDCs in the CNH (20 ss) and in the notochord (26 ss). CNH, chordoneural hinge. **c, d**, Immunostaining of Laminin (c) and Fibronectin (d) in *charon:EGFP* embryos at 12 ss and 20 ss. For 12 ss, dorsal views of KV are shown. Anterior to the top. For 20ss, lateral views of the tailbud are shown. Dorsal to the top. Anterior to the right. **e**, FISH-IF of *noto (flh)* in a *charon:EGFP* embryo at 26 ss. Lateral view. Dorsal to the top. Anterior to the left. **f**, Lineage tracing of the KVDCs cluster at the posterior tip of the CNH. H2B-mEosEM was expressed in KVDCs by *charon:mEosEM* plasmid injection into β *tin:memCherry* embryos. Photoconversion was performed in 28 ss and the labelled KVDCs were chased at 2 dpf. White arrowheads indicate the labelled KVDCs which differentiated into fin mesenchymal cells in the caudal fin fold. Scale bars, 50 µm (a-f).

The migration of KVDCs into the axial mesoderm seemed to take place exclusively in the CNH. The developing and mature notochord is surrounded by the basement membrane formed by deposition of extracellular matrix (ECM) such as Laminin and Fibronectin (Scott and Stemple, 2004). However, accumulation of these two major ECM components was reduced around the CNH as indicated by immunostaining (Fig.6d, e), which may facilitate cell mingling and infiltration of KVDCs into the CNH.

Lastly, we tracked the fate of the posterior-most KVDCs cluster which stays around the CNH until the end of the segmentation period (Fig.3d). We found that these KVDCs are distinguished from CNH cells, because they did not express the CNH markers, *noto* and (Dheen et al., 1999; Fig.6c). Using *charon:H2B-mEosEM*, we examined the fate of these KVDCs by photoconverting H2B-mEosEM-expressing cells located posterior to the CNH at around 28 ss and chased the labelled cells in 2 dpf larvae. The result demonstrated that they preferentially differentiated into fin mesenchymal cells in the caudal fin fold rather than into the CNH or notochord cells (n = 3 embryos, Fig.6f). These results imply that these KVDCs occupy a posterior-most part of the paraxial (but not the axial) mesoderm and directly differentiate into caudal fin mesenchyme.

## Discussion

During differentiation, the potency of cells becomes restricted as they are specified into certain lineages. In some contexts, however, differentiated cells can be converted to other cell types; a phenomenon called adult cell plasticity (Merrell and Stanger, 2016). Such plasticity is often observed in the regeneration process, but it is still a matter of debate whether it takes place commonly during normal development of animals (Merrell and Stanger, 2016). The present study demonstrated that derivatives of KV-epithelial cells (KVDCs) join mesodermal progenitors after KV-collapse and differentiate into mature cell types according to their location. Such “reemployment” of KV-epithelial cells is a rare example of cell plasticity of fully differentiated cells during normal development in vertebrates, which is achieved through transdifferentiation (Fig.7).

**Figure 7.**
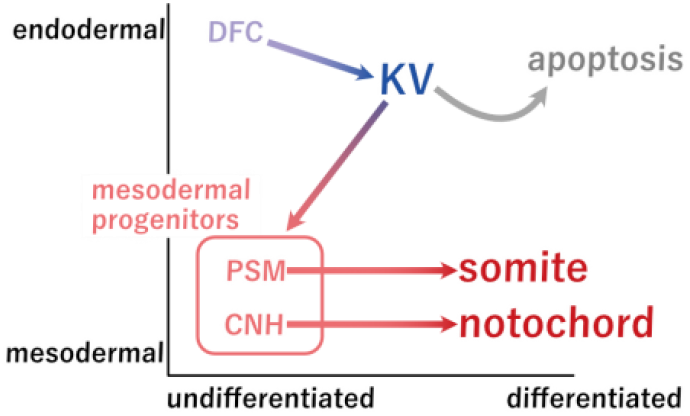
An overview of KVDCs differentiation Schematic diagram depicting the process of the transdifferentiation-mediated reemployment of KV-epithelial cells. After KV-collapse, KV-epithelial cells dedifferentiate and undergo EMT. Mesenchymalized KVDCs infiltrate into mesodermal progenitors, the PSM and CNH, and acquire new fates according to their new locations. Note that they do not directly migrate to the segmented somites or the mature notochord.

### KVDCs are a multipotent mesodermal cell population in the tailbud

Our results revealed that KVDCs in the zebrafish tailbud possess unique differentiation potency. The zebrafish tailbud contains three undifferentiated cell populations, the PSM, CNH, and neuromesodermal progenitors (NMPs; Beck, 2015; Sambasivan and Steventon, 2021). KVDCs occupy a part of the PSM and CNH, and differentiate into the paraxial and axial mesoderm. In contrast, KVDCs never contribute to NMPs as they do not give rise to the neural tube (Henrique et al., 2015). Importantly, KVDCs are unique among these progenitors in the tailbud in that they are the descendants of functional epithelial cells with primary cilia, which exerted a crucial role in left-right patterning.

The contribution of the LRO cells to the mesodermal tissues could be a common feature among vertebrates, although the origin and structures of the LRO are diverse (Blum et al., 2014). The gastrocoel roof plate in amphibians and node in mammals, which function as the LRO in these species (Blum et al., 2014), are suggested to differentiate into the notochord and somites, and into the posterior notochord, respectively (Brennan et al., 2002; Shook et al., 2004). Thus, it is likely that the vertebrate LRO cells generally behave as mesodermal cells at later stages, although DFCs in the zebrafish are transiently endodermal at the beginning of their emergence (Alexander and Stainier, 1999).

### Transdifferentiation of KVDCs

Transdifferentiation of mature cells is generally achieved through dedifferentiation into a potent state followed by redifferentiation into a new lineage (Jopling et al., 2011). The development of KVDCs appears to go through this process: KVDCs redifferentiate into somite-derivatives and notochord after being integrated into their progenitors. We observed that KV-epithelial cells undergo EMT during KV-collapse. EMT frequently occurs in association with dedifferentiation as seen in cancer metastasis and neural crest delamination (Yang and Weinberg, 2008). Indeed, EMT in KV is accompanied by the loss of differentiated epithelial characters such as primary cilia and tight junctions, which lead to KV-collapse and cell migration. Factors triggering this EMT are yet to be determined, although potent signalling ligands, Wnt, BMP and FGF are produced in the nearby PSM (Hubaud and Pourquié, 2014; Row and Kimelman, 2009). Furthermore, we described the shrinkage of the YSL underlying KV in the present study (Fig.4c). This shrinkage might generate mechanical force which triggers EMT in cooperation with those signalling ligands (Gjorevski et al., 2012).

In general, mature epithelial cells maintain their fate and integrity by cell-cell junctions and attachment to the basement membrane (Yang et al., 2020). Then, why can KV-epithelial cells undergo transdifferentiation and what makes them so unique? Despite the presence of evident apico-basal polarity and cell-cell junctions, KV epithelium seems to lack the basement membrane, as revealed by our immunostaining and a previous report on the atypical localization of Laminin β1a to their apical side (Hochgreb-Hägele et al., 2013). Thus, KV-epithelial cells may not be of typical epithelium in character, which could reflect their transient function. Lack of the basement membrane would allow KV-epithelial cells to dedifferentiate and migrate without a need of ECM remodelling.

Some vertebrate tissues consist of cells derived from different lineages. The prominent examples are the zebrafish pituitary and mammalian vascular endothelium, where undifferentiated cells from different origins converge at the transcription level once they are intermingled (Fabian et al., 2020; Plein et al., 2018). We confirmed here that KVDCs, whose origin is distinct from the authentic mesoderm, differentiate into the notochord and somite-derivatives in the caudal region. KVDCs are thus a good model for the study of such lineage convergence during normal development. Single-cell multiomics using *charon* transgenic lines will provide insights into the mechanism of how and to what extent cells from different lineages converge together to form a functionally integrated tissue.

## Figure legends

**Supplementary Figure 1.**
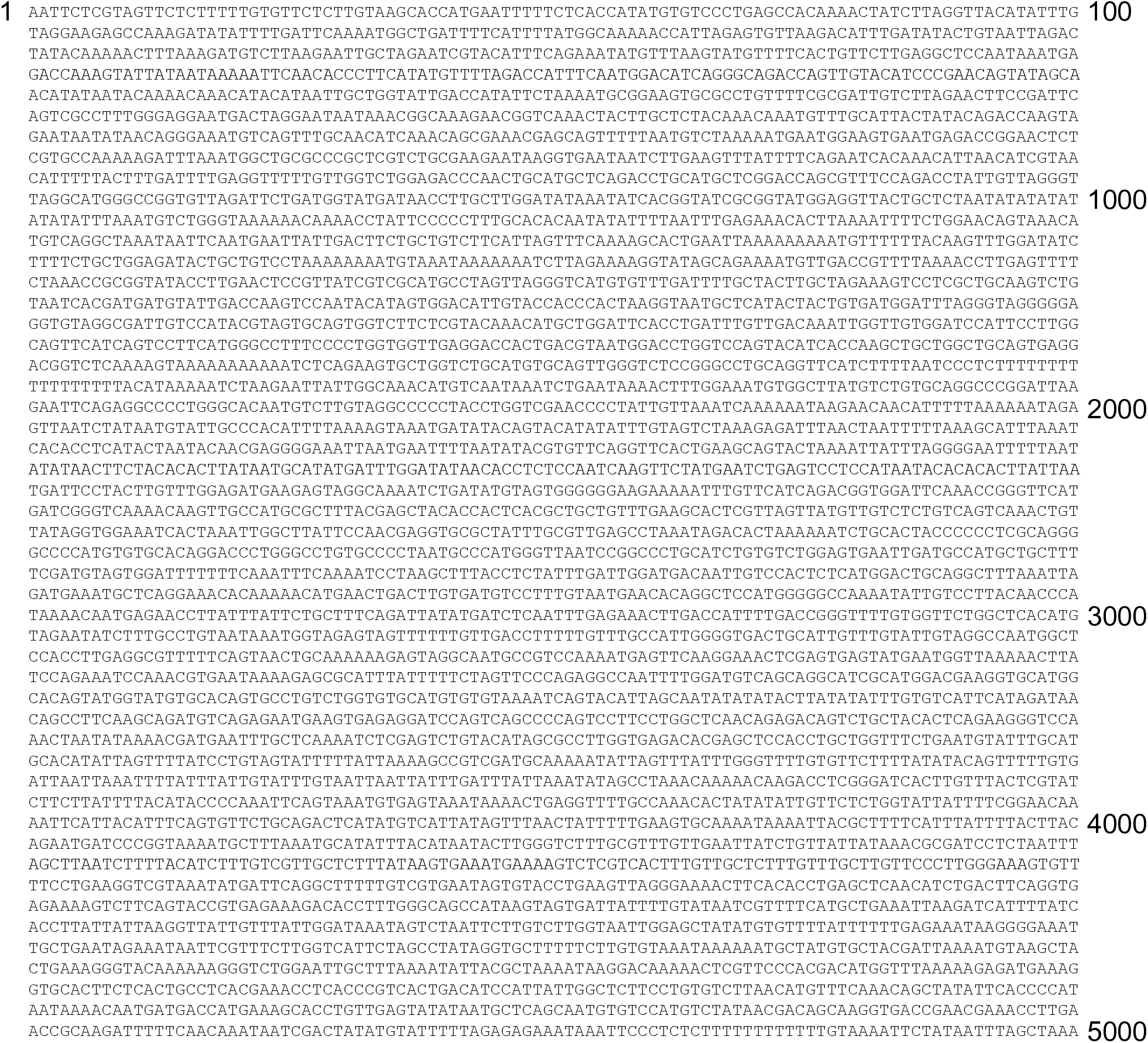
Sequence of the zebrafish 5-kb *charon* promoter Nucleotide sequence of the 5-kb *charon* promoter in the zebrafish.

**Supplementary Figure 2.**
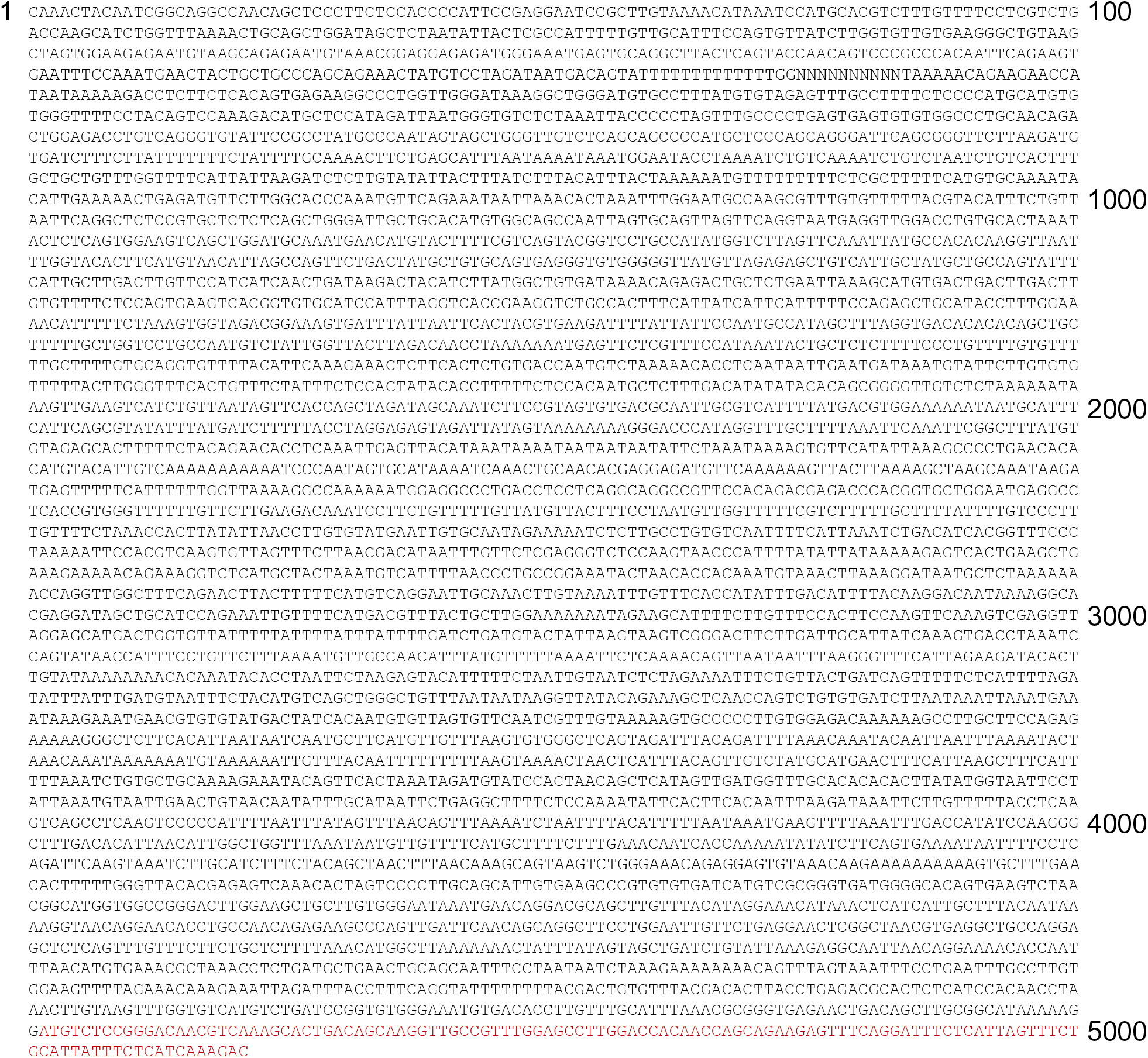
Sequence of the medaka 5-kb *charon* promoter Nucleotide sequence of the 5-kb *charon* promoter in medaka. Red, partial coding sequences of *charon* gene.

**Supplementary Figure 3.**
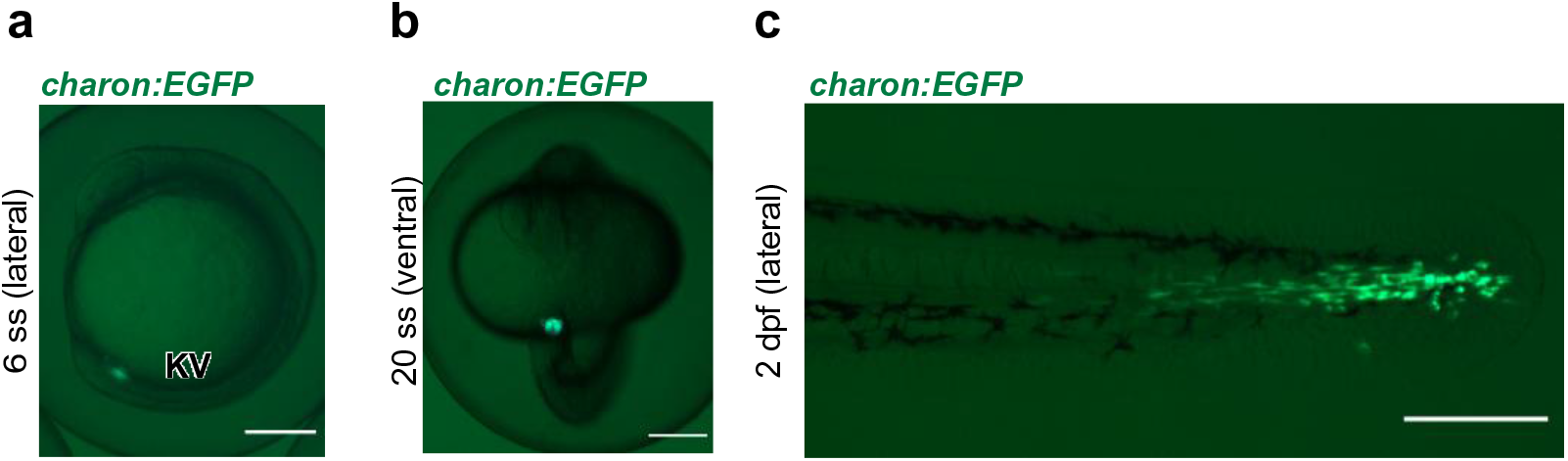
Fluorescence stereomicroscope images of *charon:EGFP* **a**, Lateral view of a *charon:EGFP* embryo at 6 ss. **b**, Ventral view of a *charon:EGFP* embryo at 20 ss. **c**, Lateral view of a *charon:EGFP* larva at 2 dpf. Scale bars, 200 µm (a-c).

**Supplementary Figure 4.**
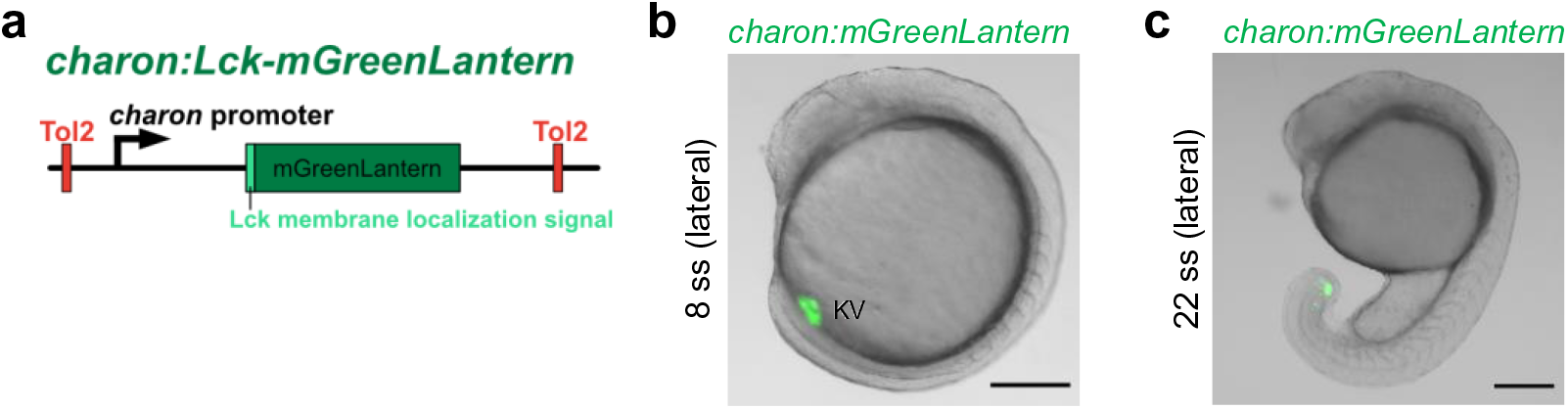
Fluorescence of *charon:Lck-mGreenLantern* **a**, The construction of *charon:Lck-mGreenLantern* transgene. **b, c**, Lateral view of a *charon:Lck-mGreenLantern* embryo at 8 ss (b) and 22 ss (c). KV, Kupffer’s vesicle. Scale bars, 200 µm.

**Supplementary Figure 5.**
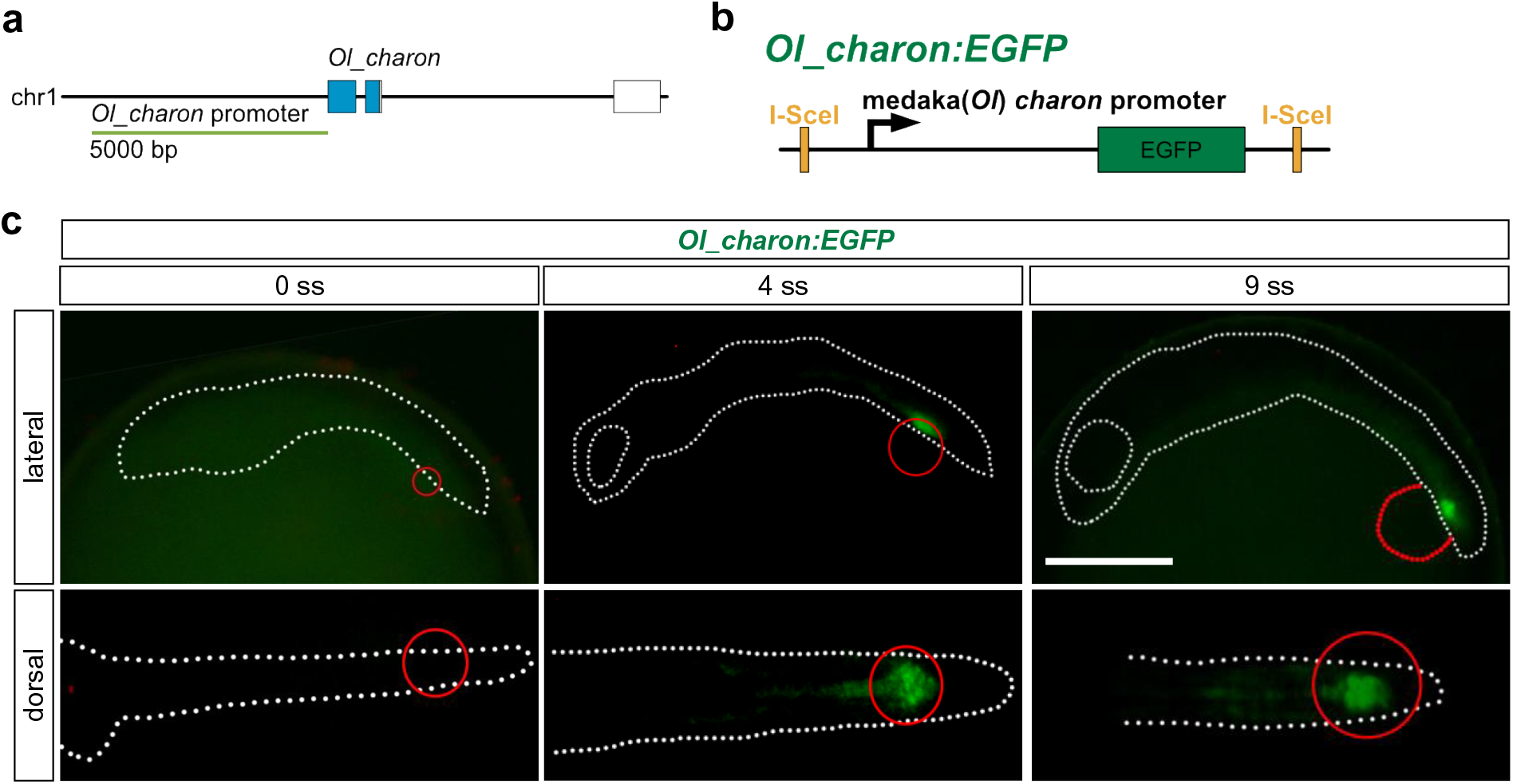
Medaka *charon* promoter is active in medaka KV-epithelial cells **a**, Location of the *charon* 5-kb promoter in the medaka genome. *charon* resides on chromosome 1, and the *charon* 5-kb promoter spans from the direct upstream position of its start codon. **b**, Construction of *Ol_charon:EGFP* transgene flanked by I-SceI recognition sites. **c**, Expression of EGFP in *Ol_charon:EGFP* embryos. Lateral views on the top row, dorsal views on the bottom row. White dot lines indicate embryonic bodies. Red circle indicates the position of KV. Scale bars, 500 µm.

**Supplementary Figure 6.**
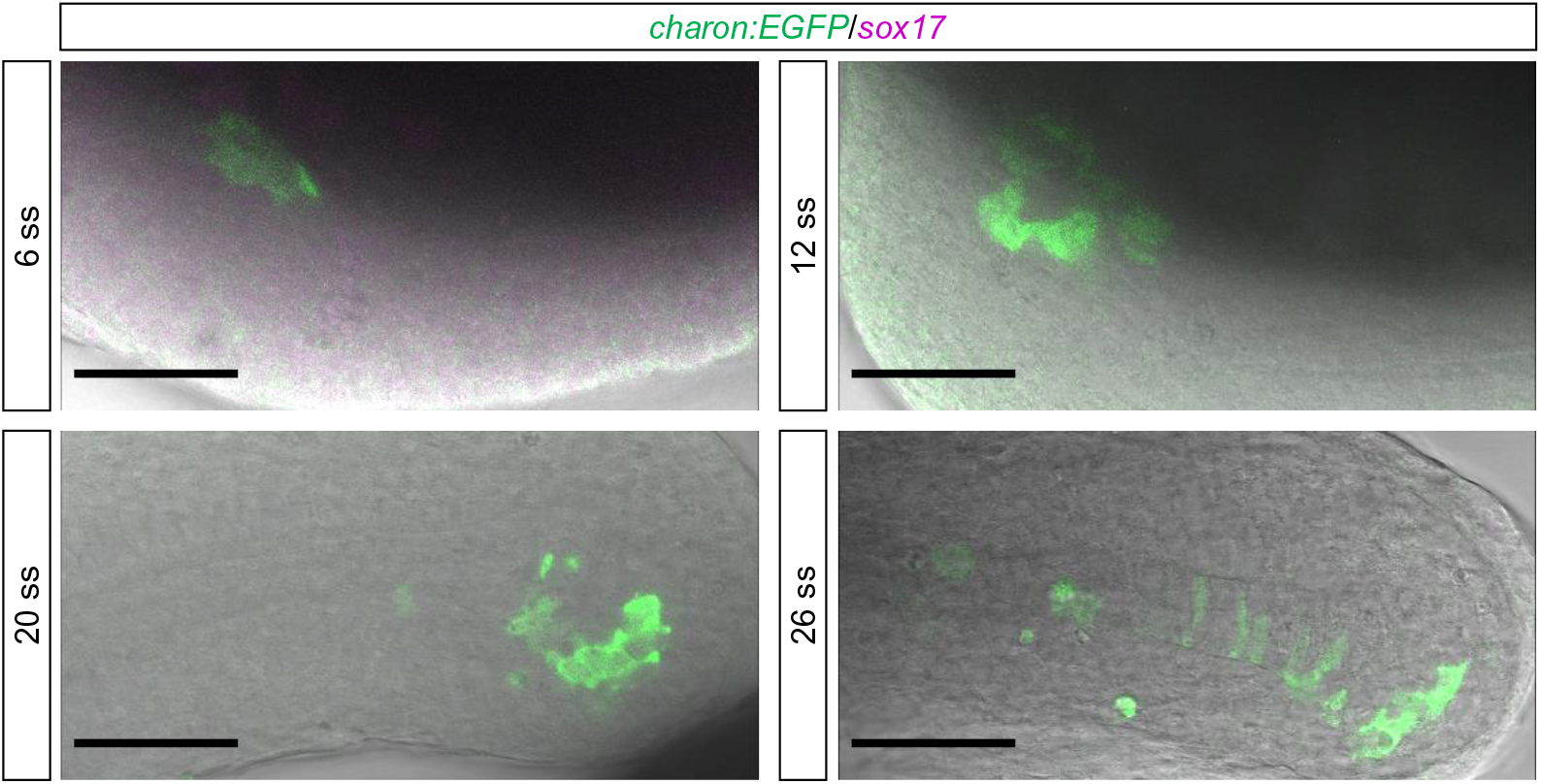
FISH-IF of *sox17* in *charon:EGFP* embryos FISH-IF of *sox17* in *charon:EGFP* embryos at 6 ss, 12 ss, 20 ss and 26 ss. Lateral views of tailbud regions are shown. Scale bars, 50 µm.

**Supplementary Figure 7.**
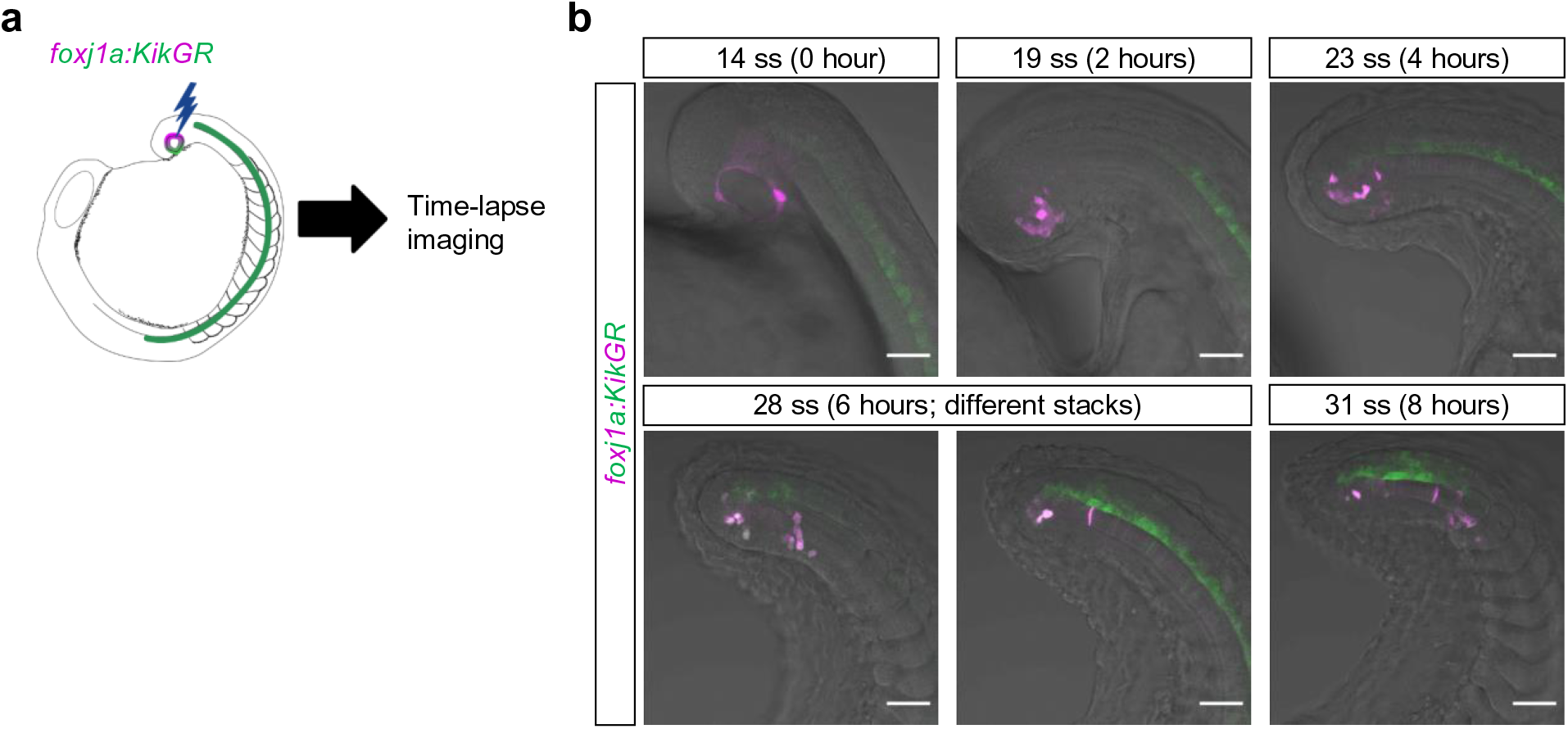
*foxj1a:KikGR* supports KV-epithelial cell migration **a**, Scheme for the photoconversion labelling of KV-epithelial cells in *foxj1a:KikGR*. KikGR in KV-epithelial cells was selectively photoconverted at around 13 ss and time-lapse images were taken every 1 hour. **b**, Time-lapse images of a photoconverted *foxj1a:KikGR* embryo. Single stacks from images taken every 2 hours after photoconversion are shown. When photoconverted at around 13 ss, labelled KVDCs migrated into the PSM and the notochord by 28 ss (6 hours after photoconversion). Scale bars, 50 µm.

**Supplementary Figure 8.**
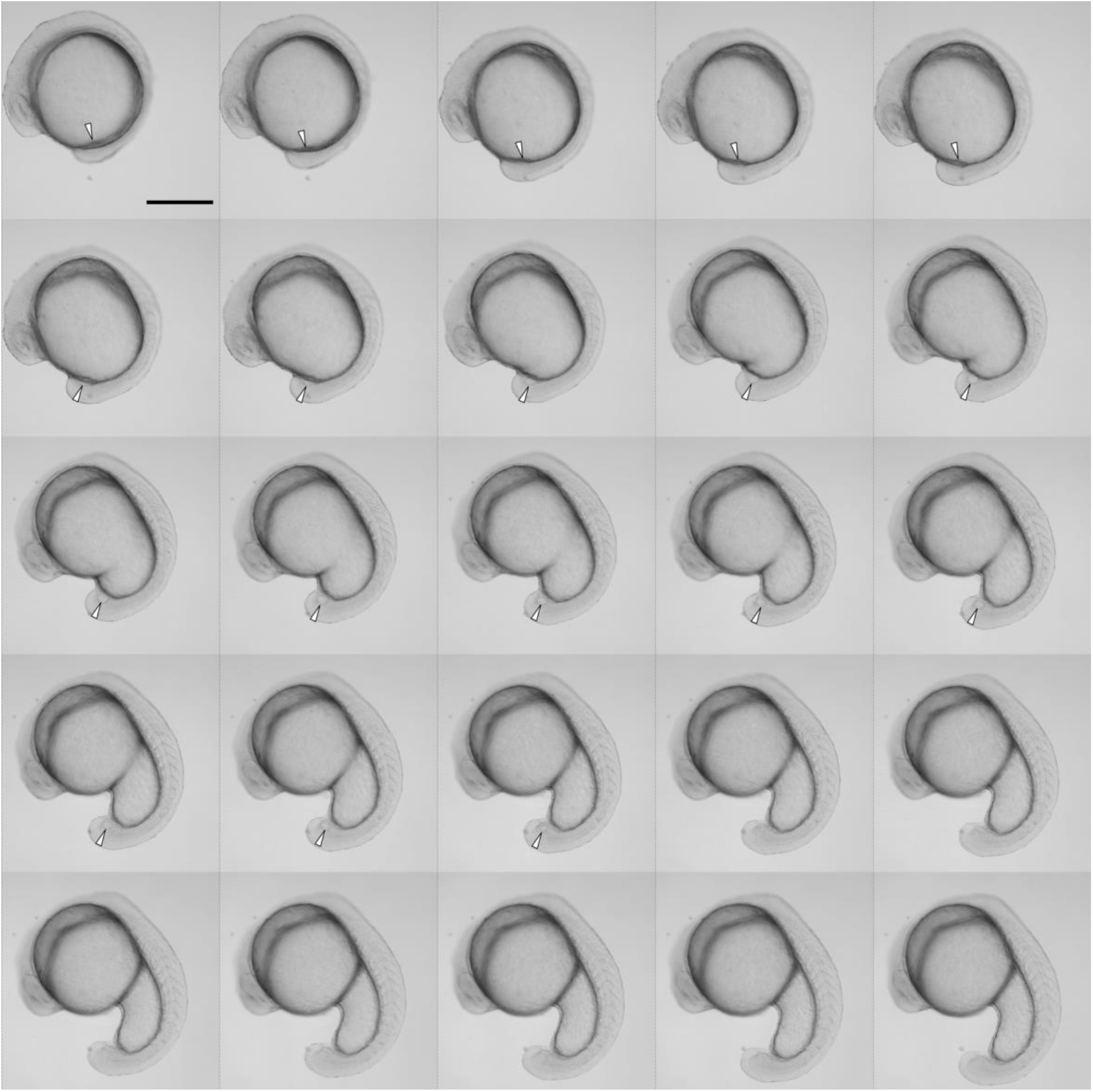
Time-lapse imaging of the incorporation of KV into the tailbud Time-lapse imaging of the KV development in a TL2E embryo. Arrowheads indicate the position of KV. Staring from 4 ss, images are taken every 15 minutes for 10 hours. Scale bars, 50 µm.

Supplementary Video 1. Time-lapse imaging of KV-collapse

A video showing the process of KV-collapse in a *charon:EGFP;*β*actin:memCherry* embryo. Time-lapse images were taken every 5 minutes for 3 hours from 13 ss. Note that a ring-like structure is formed beneath KV during KV-collapse. Scale bar, 50 µm.

Supplementary Video 2. Formation of filopodia-like structures in KVDCs

Green channel is extracted from Supplementary Video 1 to show the formation of filopodia-like structure in KVDCs. Scale bar, 50 µm.

## Author Contributions

Conceptualization, T.I., K.I., and H.T.; Methodology, T.I., K.I., and T.K.; Investigation, T.I., K.I., and T.K.; Resources, T.I., K.I., and T.K.; Writing - Original Draft, T.I.; Writing - Review & Editing, T.I., T.K., and H.T.; Supervision, H.T.; Funding Acquisition, T.I., K.I., T.K. and H.T..

## Acknowledgement

We thank S. Megason for β*actin:memCherry* strain; C. Mosimann for *drl:mCherry* strain and *drl:EGFP (pCM298)* plasmid; N. H. Patel for Pax3/7 antibody (DP312); A. Shimada for technical support and discussion; Y. Yamagishi, S. Tayama, I. Fukuda, M. Funato and M. Sakamoto for fish husbandry. This work was supported by Grant-in-Aid for Japan Society for the Promotion of Science (JSPS) Fellows under Grant Numbers 18J21960 (T.I.) and 15J08599 (K.I.), JSPS KAKENHI under Grant Numbers JP19K23741 and JP21K15101 (T.K.) and Japan Agency for Medical Research and Development (AMED) under Grant Number JP21gm1110007 (H.T.).

## Material and Methods

### Zebrafish strain and manipulation of embryos

The RW (RIKEN WT) and TL2E (Tüpfel long fin 2E) strains were used as the wild-type zebrafish strains. Adult fish and embryos were maintained under standard conditions. Fertilized embryos were incubated at 23-28°C in 1/3×Ringer’s solution (38.7 mM NaCl, 0.97 mM KCl, 1.67 mM HEPES, 1.80 mM CaCl_2_) to obtain the stage of interest. For imaging hatched larvae, 0.003% N-phenylthiourea (PTU) was added to 1/3×Ringer’s solution during somitogenesis to prevent pigmentation. Staging of embryos is based on Kimmel et al. (1995). All experimental procedures and animal care were carried out according to the animal ethics committee of the University of Tokyo (Approval No. 20-2).

### Plasmid construction

The sequences of *charon* and *foxj1a* promoters (5 kb and 5.2 kb, respectively) were amplified from the genomic DNA of the RW strain and were cloned into pCR Blunt II-TOPO vector (Invitrogen). pDestTol2pA2*-charon:EGFP* (*charon:EGFP* for short) was constructed by replacing the *drl* promoter in pDestTol2pA2*-drl:EGFP* (Mosimann et al., 2015) with the cloned *charon* promoter. Medaka *charon* promoter was cloned from genomic DNA of medaka d-rR strain and was inserted into pBlueScript II SK(+)-EGFP carrying an I-SceI cassette to make pBSSK-I-SceI-*Ol_charon:EGFP* (*Ol_charon:EGFP* for short). pDestTol2pA2-*charon:Lck-mGreenLantern* was constructed as follows: the CDS of mGreenLantern (Campbell et al., 2020) whose codon usage is optimized for zebrafish was purchased from Integrated DNA Technologies (IDT), and was inserted into pCSf107mT vector (Mii et al., 2009) with an N-terminal membrane localization signal of mouse Lck (Chertkova et al., 2017) to make pCSf107-Lck-mGreenLantern. Then, the CDS of Lck-mGreenLantern was subcloned into pDestTol2pA2 vector carrying the *charon* promoter. pDestTol2pA2-*charon:H2B-mEosEM* was constructed as follows: the CDS of mEosEM was synthesized through mutagenesis of mEos4b (Paez-Segala et al., 2015) as described (Fu et al., 2020), and was assembled together with the CDS of human Histone H2B type 1-J (H2BC11) into pCSf107mT to make pCSf107-H2B-mEosEM. Then, the CDS of H2B-mEosEM was subcloned into pDestTol2pA2 vector carrying the *charon* promoter. To construct template plasmids for the antisense probe synthesis, full-length coding sequences (CDSs) of zebrafish *charon*, *sox17*, *msgn1*, *myod1*, *nkx3.1*, *ntl* and *noto* cloned from 16 ss cDNA were inserted into pCSf107mT. These cloning and subcloning were performed using PrimeSTAR GXL polymerase, PrimeSTAR MAX polymerase, In-fusion HD kit and In-Fusion Snap Assembly kit (TaKaRa). pRSETa_mEos4b was a gift from Loren Looger (Addgene plasmid #51073; http://n2t.net/addgene:51073; RRID:Addgene_51073). Lck-mScarlet-I was a gift from Dorus Gadella (Addgene plasmid #98821; http://n2t.net/addgene:98821; RRID:Addgene_98821). Digital plasmid maps are available upon request.

### Transgenesis

Zebrafish transgenesis was based on the Tol2 transposase method (Kikuta and Kawakami, 2009). To obtain founders, Tol2 mRNA (25 ng/µL) and plasmids (12.5 ng/µL) were injected into one-cell stage embryos of the RW strain. Founders carrying transgenes were screened by crossing with the TL2E strain. For medaka transgenesis, the I-SceI method (Rembold et al., 2006) was used. Following digestion with I-SceI, *Ol_charon:EGFP* plasmid was injected into one-cell stage embryos of the d-rR wildtype strain. Founders were screened by crossing with the d-rR strain. Fluorescence image of transgenic embryos was captured by M165 FC fluorescence stereomicroscope and DFC7000T digital camera (Leica).

### Time-lapse imaging and photoconversion

To perform time-lapse imaging, embryos were mounted laterally in the chamber of 1% agarose in 1/3×Ringer’s solution with their left side down. Embryos were imaged by LSM710 confocal microscopy (Carl Zeiss) from 13 ss for 3 hours or longer at an interval of 5 min. In cell-tracking experiments, the photoconversion of H2B-mEosEM was performed with a 405 nm diode laser at the power of 10%. After photoconversion, embryos were released in 1/3x Ringer’s solution with 0.003% PTU and were grown until they reach the stages of interest in a shaded chamber. The 3D rendering of time-lapse images was performed with Imaris (version 8.1.2, Bitplane).

### Histology and immunostaining

Embryos were fixed with 4% paraformaldehyde (PFA) in PBST at 4°C overnight or at room temperature for 2 hours, washed with PBST for three times and were stored at 4°C until use (except for Laminin and Fibronectin, see below). Fixed embryos were permeabilized with 1% Triton X-100 in PBS, blocked with 2% BSA in PBSDT (PBS/1% DMSO/0.1% Triton X-100) and were immunostained with following antibodies: mouse monoclonal anti-GFP (1/100, A-11120, Invitrogen), rabbit poly-clonal anti-GFP (1/800, 632592, Clontech), anti-Myosin heavy chain (1/10, F59, deposited to the DSHB by Stockdale, F.E.), anti-Myosin light chain 1 slow and 3 fast (1/10, F310, deposited to the DSHB by Stockdale, F.E.), anti-Pax3/7 (1/500, DP312, bycourtesy of N. H. Patel), anti-DsRed (1/500, 632496, Clontech), anti-cleaved caspase-3 (1/500, 9661, Cell Signaling Technology), anti-phospho Histone H3 (1/500, 06-570, Merck), and anti-ZO-1 (1/50, 33-9100, Invitrogen). For immunostaining of Laminin and Fibronectin, embryos were fixed by 2% PFA for 2 hours at 25°C, then for overnight at 4°C. After quenching PFA with 50 mM glycine in PBSTT (PBS/0.5% Triton X-100/0.5% Tween 20) and subsequent washing, embryos were blocked with 5% BSA in PBSTT, and were stained with anti-Laminin (1/100, L9393, Sigma-Aldrich) or anti-Fibronectin (1/100, F3648, Sigma-Aldrich). Secondary antibodies for immunofluorescence were anti-rabbit IgG labelled with Alexa Fluor 488, 555, or 568 (A11008, A21429 or A10042, Invitrogen) and anti-mouse IgG labelled with Alexa Fluor 488 or 568 (A11001 or A10037, Invitrogen). For counterstaining, Phalloidin (labelled with Rhodamine or Alexa Fluor 647) (R415 or A22287, Invitrogen) and DAPI (4’,6-diamidino-2-phenylindole) were added to the secondary antibody solution. Stained embryos were mounted with 1% LMP agarose/PBS (16520-050, Invitrogen) in a glass-based dish (3911-035, Iwaki) and were imaged by LSM710 confocal microscopy (Carl Zeiss). 40× and 25× water-immersion objective lenses (LD C-Apochromat 40x/1.1 W Korr M27 and LD LCI Plan-Apochromat 25x/0.8 Imm Corr DIC M27, Carl Zeiss) and a 5× objective lens (EC Plan-Neofluar 5x/0.16) were used. The contrast adjustment and cropping of captured images were performed by Fiji (version 1.53c; Schindelin et al., 2012).

For immunostaining of cross sections in 3 dpf larvae of *charon:EGFP*, fixed specimens were embedded in 4% LMP agarose/PBS, and 100-200 µm sections were prepared using Vibratome 3000 Tissue Sectioning System. Free-floating sections were stained with appropriate antibodies and mounted with 50% Glycerol/PBS on a slide glass.

For DAB staining of EGFP-expressing cells, fixed *charon:EGFP* embryos were blocked with 0.3% H_2_O_2_ for 30 min and with 2% BSA in PBST for 2 hours, stained with anti-GFP antibody (1/800, 632592, Clontech) and biotinylated anti-rabbit IgG (1/200, BA-1000, Vector Laboratories). VECTASTAIN® Elite® ABC Kit (Vector Laboratories) and DAB tablets (D5905, Sigma-Aldrich) were used for DAB chromogenic reaction. Stained embryos were embedded in Technovit 7100 resin (Kulzer Technique) and 5 µm sections were obtained using an RM2245 microtome (Leica). Sections were counterstained with Mayer’s hematoxylin solution and images were captured by BX61 microscope (Olympus) and AxioCam MRc 5 camera (Carl Zeiss) using a 40× objective lens.

### Fluorescence *in situ* hybridization and immunofluorescence (FISH-IF)

*charon:EGFP* embryos at stages of interest were processed according to He et al., (2020) with minor modifications. Briefly, fixed embryos were permeabilized with 1% Triton X-100 in PBS for 1 hour and were hybridized with DIG-labelled probes at 60°C. Chromogenic reaction was performed with either TSA Plus Cyanine 3 Kit (Akoya Biosciences) for *nkx3.1* or ImmPACT Vector Red (Vector Laboratories) for the other genes. Afterward, embryos were subjected to immunostaining against EGFP using anti-GFP antibody (1/500, A11122, Invitrogen).

